# Survival Analysis on Rare Events Using Group-Regularized Multi-Response Cox Regression

**DOI:** 10.1101/2020.06.21.163675

**Authors:** Ruilin Li, Yosuke Tanigawa, Johanne M. Justesen, Jonathan Taylor, Trevor Hastie, Robert Tibshirani, Manuel A. Rivas

## Abstract

We propose a Sparse-Group regularized Cox regression method to improve the prediction performance of large-scale and high-dimensional survival data with few observed events. Our approach is applicable when there is one or more other survival responses that 1. has a large number of observed events; 2. share a common set of associated predictors with the rare event response. This scenario is common in the UK Biobank (Sudlow et al. 2015) dataset where records for a large number of common and rare diseases of the same set of individuals are available. By analyzing these responses together, we hope to achieve higher prediction performance than when they are analyzed individually. To make this approach practical for large-scale data, we developed an accelerated proximal gradient optimization algorithm as well as a screening procedure inspired by Qian et al. (2019). We provide a software implementation of the proposed method and demonstrate its efficacy through simulations and applications to UK Biobank data.

## 1 Introduction

### 1.1 Cox Proportional Hazard Model

Cox model (Cox 1972) provides a flexible mathematical framework that describes the relationship between the predictors and a time-to-event response. For each individual we observe a triple {*O, X, T*}, where *X* ∈ ℝ^*d*^ are the features and *O* ∈ 0, 1 is a status indicator. If *O* = 1, then *T* is the actual time-to-event. If *O* = 0, then we only know that the time-to-event is at least *T*. The hazard function according to the Cox model can be written as

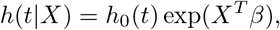

where *β* ∈ ℝ^*d*^ is the coefficients vector that measures the strength of association between *X* and the response. This hazard function is equivalent to the cumulative distribution function:

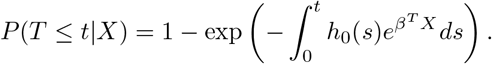

Here *h*_0_: ℝ^+^ ↦ ℝ^+^ is the baseline hazard function. In our applications we are interested in the relationship between the features and the responses, so the baseline hazard function is a nuisance variable. We can estimate the parameters *β* directly without knowing the baseline hazard by maximizing the log partial likelihood function (Cox 1972):

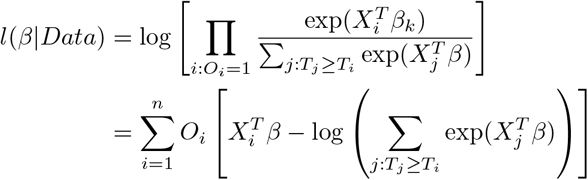

When the number of observed events is small relative to *n*, estimating *β* becomes challenging. This could happen, for example, when the time-to-event response is the age of diagnosis of a rare disease. In particular, if *O*_*i*_ are i.i.d Bernoulli random variables with probability *p*, then the information matrix is proportional to *p* and thus the asymptotic variance of the maximum partial likelihood estimate is inversely proportional to *p*.

We evaluate a fitted survival model using the concordance index, or the C-index. For a parameter estimate 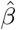, the C-index is defined as

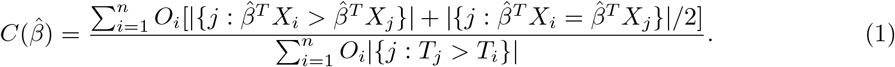

For more details on C-index, see Harrell et al. (1982), Li et al. (2020).

### 1.2 Sparse-Group Lasso

The Lasso method (Tibshirani 1996) makes the assumption that only a small subset of predictors are associated with the response. In other words, it assumes that *β* has only a small number of non-zero entries. A sparse solution can be obtained by optimizing an *L*_1_-regularized objective function.

Sparse-Group Lasso (Simon et al. 2013) assumes not only that many individual elements of *β* are 0, but also that many groups of variables have coefficients 0 simultaneously. For example, in a single-response Cox model with *d*-dimensional features, if groups of variables 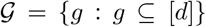 are believed to have sparse-group structure, then the sparse-group Lasso minimizes the following objective function:

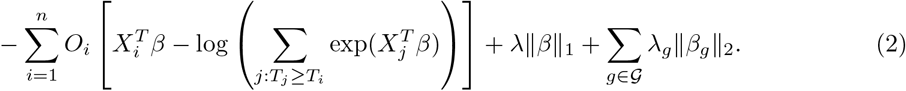

## 2 Methods

### 2.1 Preliminaries

In this section we define the notations and the key model assumptions that we will use in the subsequent sections. For an integer *n*, define [*n*] = {1, 2, · · · *, n*} and define *x*_+_ = max{*x,* 0} for all *x* ∈ ℝ.

We analyze *K* ≥ 1 time-to-event responses on *n* individuals. For example, the responses could be time from birth to *K* different diseases. The data we observed are in the format:

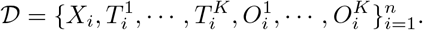

Here *X*_*i*_ ∈ ℝ^*d*^ are *i*th individual’s features. Denote the full features matrix ***X*** = [*X*_1_, *X*_2_, · · ·, *X*_*n*_]^*T*^ ∈ ℝ^*x*×*d*^. For *k* = 1, · · ·, *K*, 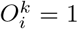 implies that 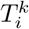 is the true time until the event *k* and for the *i*th individual, and 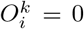 implies that the true time until the event *k* is right-censored at 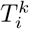. We assume each response follows a Cox proportional hazard model:

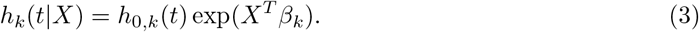

where *h*_0*,k*_: ℝ^+^ ⟼ ℝ^+^ is the baseline hazard function of the *k*th response. Let

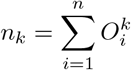

be the number of observed event *k*.

We make the assumption that not only *β*_*k*_ is sparse for all *k* ∈ [*K*], but they also have a small common support. That is 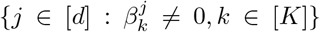 is a small set relative to *d*. In human genetics applications, the first assumption means that each response is associated with a small set of genetic variants, and the second assumption implies that there are significant overlap among the genetic variants that are associated with each response. This belief is the main driver for prediction performance improvements on rare diseases, and it translates to the following regularized partial likelihood objective function:

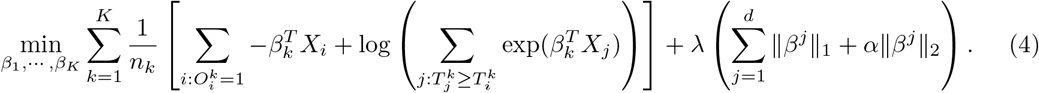

Here the first term is the sum of the *K* negative log-partial-likelihood, normalized by the number of observed events. The second term is the regularization. ||*β*^*j*^||_1_, ||*β*^*j*^||_2_ are the 1-norm and 2-norm of the coefficients for the *j*th variable. That is, if we put all the parameters into a matrix *B* = [*β*_1_, *β*_2_, · · ·, *β*_*K*_] ∈ ℝ^*d×K*^, then *β*^*j*^ is the *j*th row of *B* and *β*_*k*_ is the *k*th column. Note that when *α* = 0, the objective function decouples for each *β_k_* and they can be optimized separately. In our implementation we solve a slightly more general problem:

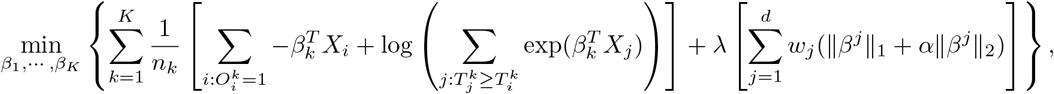

where 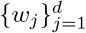 are user provided penalty factor for each variables, which may be useful in the setting where protein-truncating or protein-altering variants should be prioritized (Rivas et al. 2015, DeBoever et al. 2018). Just like in Yuan & Lin (2006), our implementation by default fixes *α* at 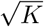. The solution is computed on a pre-defined sequence of *λ*s: *λ*_1_ > *λ*_2_ > … > *λ*_*L*_, where *λ*_1_ is chosen so that the solution just become non-zero.

To simplify the notation, for *j* ∈ [*d*] denote

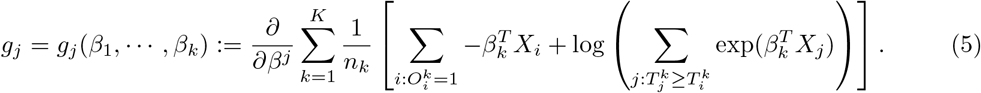

Here *g*_*j*_ ∈ ℝ^*K*^ is the partial derivative of the smooth part of (4) with respect to the coefficients of the variable *j*. Finally, let *S*_1_(·; *λ*): ℝ^*K*^ ⟼ ℝ^*K*^ be the element-wise soft-thresholding function:

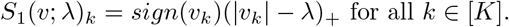

### 2.2 Optimization Method

We use a Nesterov-accelerated (Nesterov 1983) proximal gradient method (Daubechies et al. 2004, Beck & Teboulle 2009) to optimize the objective function (4). Proximal gradient algorithm is particularly suitable when the objective function is the sum of a smooth function and a simple function. In our case the smooth function is the sum of the negative log-partial-likelihood functions, and the simple function is the regularization term. The algorithm alternates between two steps until convergence criteria is met:

1. A gradient step that decreases the smooth part of the objective:

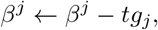

where *t* is the step size that we determine using backtracking line search.
2. A proximal step that keeps the regularization term small:

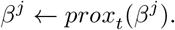 Here the proximal operator *prox*_*t*_: ℝ^*K*^ ⟼ ℝ^*K*^ is defined as

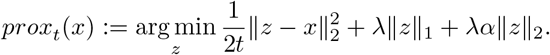 To simplify the notation we omit the dependency of *prox*_*t*_(*x*) on *λ, α*. Simple calculation shows that

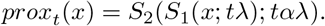

where *S*_2_(·; *tαλ*): ℝ^*K*^ ⟼ ℝ^*K*^ is a group soft-thresholding operator:

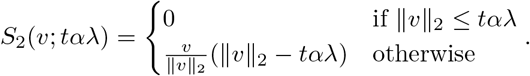

We describe the optimization algorithm in pseudocode, including details about Nesterov acceleration and backtracking line search in algorithm 1.

### 2.3 Variable Screening for Lasso Path

In many of our applications the data is large-scale and high-dimensional. For example, the UK Biobank dataset (Sudlow et al. 2015) contains millions of genetic variants and over 500, 000 participants. Reading the feature matrix of the UK Biobank dataset into R requires more than 4 terabytes of memory, which is much larger than the RAM size of most computers. While memory mapping (Kane et al. 2013) allows users to access out-of-memory data with ease, it requires lots of disk Input/Output operations, which is much slower than in-memory operation. This becomes even more problematic for iterative optimization algorithms that use the entire feature matrix every iteration.

To reduce the frequency of reading the full data matrix, here we derive a version of variable screening method following similar ideas of the strong rule (Tibshirani et al. 2012) and the Batch Screening Iterative Lasso (Qian et al. 2019).

For each *α* > 0, *v* ∈ ℝ^*K*^, define the *α*^*^ norm of *v* to be

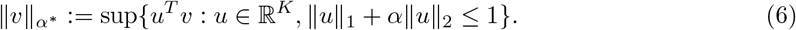

We use the following results to get an optimality condition for the solutions *β*_1_,, *β*_*K*_. The proofs are given in section 6.

**Algorithm 1:**
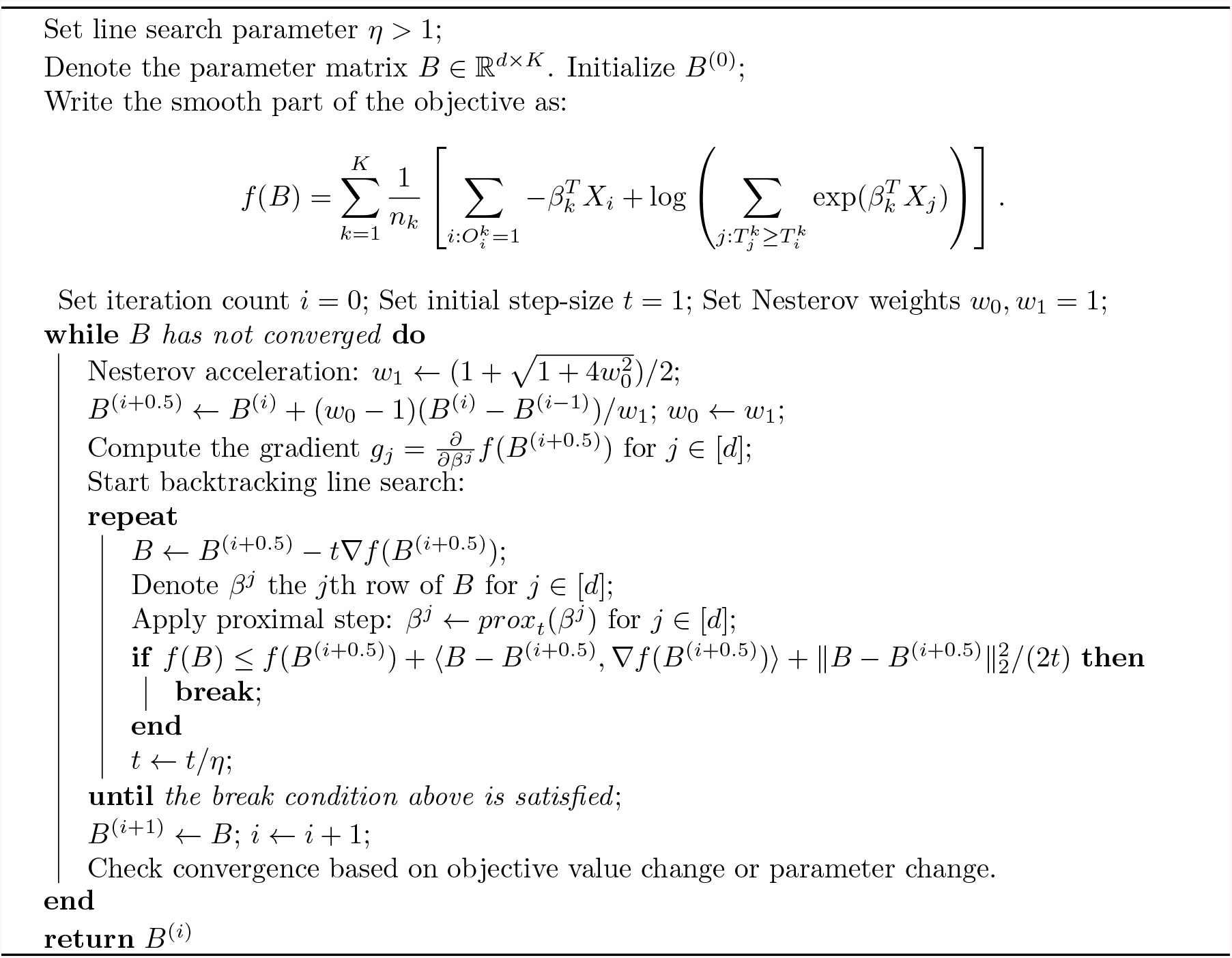
Proximal Gradient Method for (4)

#### Proposition 1.

*For any* λ > 0, *the gradients defined at* (5) *at the optimal solution to* (4) *satisfies:*

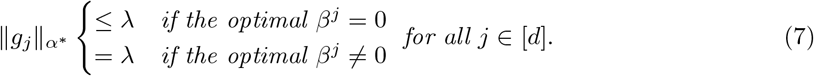

This result motivates us to first fit a model (solving (4)) using a small number of variables whose gradient has the largest *α*^*^ norm, assuming the coefficients for the rest of the variables are all zero. Then to verify the validity of the assumption we check that ||*g*_*j*_||_*α*_* ≤ *λ* for variables assumed to have zero coefficients. We refer to this step as KKT checking. Note that based on its definition (6), it’s not clear how we can compute ||*v*||_*α*_*. Here we give a more explicit characterization.

#### Proposition 2.

||*v*||_*α*_* ≤ *λ* if and only if ||*S*_1_(*v*; *λ*)||_2_ ≤ *αλ.*

Using the above we can check if ||*v*||_*α*_* ≤ *λ* quite easily and compute ||*v*||_*α*_* using binary search in 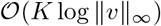 to any fixed precision (since ||*v*||_*α*_* ≤ ||*v*||_*∞*_).

Now we are ready to state the overall structure of our algorithm with variable screening. Suppose valid solutions for *λ*_1_, · · ·, *λ*_*l*_ have been obtained. Next we follow these steps:

1. (**Screening**) In the last iteration, we cache the full gradient 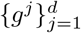 evaluated at the solution at *λ*_*l*_. In the fitting step we include two types of variables:

- We include the ever-active variables 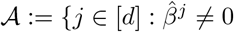 for any previously obtained 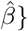}.
- Top *M* variables with the largest ||*g*_*j*_||_*α\**_ that are also not ever-active. We denote the set of variables used to fit (4) as the strong set 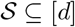.
2. (**Fitting**) In this step, we solve the problem (4) for the next few *λ*s using only variables in 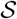, assuming *β*^*j*^ = 0 for all 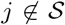. This is done using proximal gradient descent (algorithm 1). To speed-up the computation we initialize the variables at the previous valid solution (warm start).
3. (**KKT Checking**) After obtaining the solution from the fitting step. We compute the full gradient. This is the only step we will need the full data matrix ***X***. We check if the KKT conditions (7) are satisfied for all variables then go back to the screening step at the first *λ* value where the KKT condition fails. We also cache the full gradient at the last valid solutions for the screening step.
4. (**Early Stopping**) We keep a separate validation set to evaluate the current estimated parameters. We choose the optimal *λ* as the one that gives the highest validation C-index. The optimal *λ* might be different for different responses. In this paper we focus only on the prediction accuracy of one rare event, so it is reasonable to stop when the validation C-index of this response starts to decrease, regardless if the optimal *λ* for other responses has been reached.

These steps are described in algorithm 2.

### 2.4 Software

We implemented the proximal gradient method in section 2.1 as an R package, available at https://github.com/rivas-lab/multisnpnet-Cox. In this package we also implement the screening procedure described in this section for genetics data in Plink2 format (Chang et al. 2014).

**Algorithm 2:**
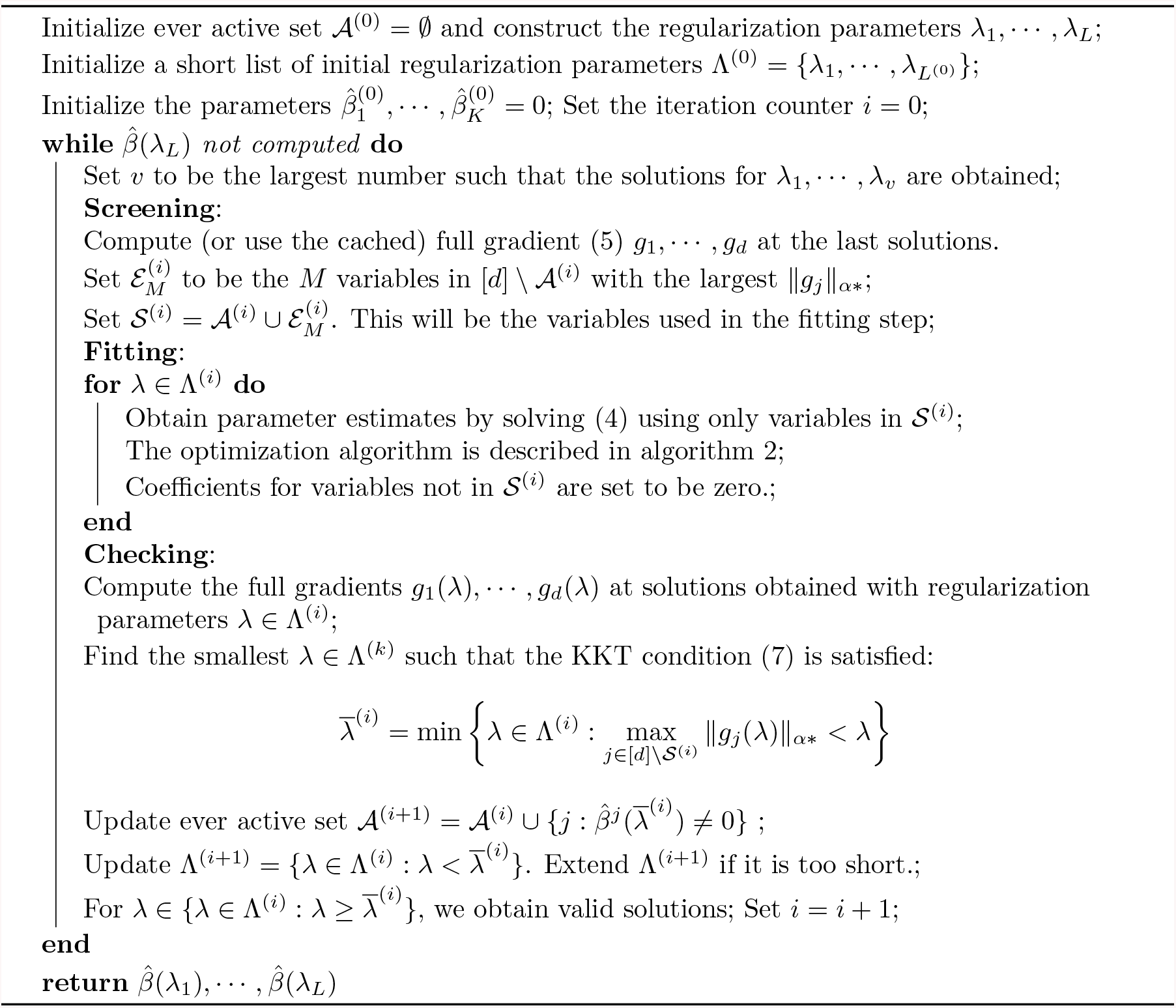
Iterative Screening for Lasso Path

The major computational bottleneck in the proximal gradient method are matrix-matrix multiplications. These operations have high arithmetic intensity and are particularly suitable for GPU acceleration. Therefore we also provide a GPU implementation of algorithm 1 on CUDA-enabled device, available at https://github.com/rivas-lab/multisnpnet-Cox_gpu. With *n* = *d* = 10000, *K* = 20, the GPU implementation achieves almost 10× speedup in solving a path of 50 *λ*s (9.7 seconds vs 92 seconds) on a Tesla V100 GPU than the CPU implementation on an Intel Xeon 6528R processor with 28 threads, accelerated using Intel’s Math Kernel Library.

## 3 Simulations

In this section we compare the performance of the proposed approach against a simple Lasso, where multiple responses are fitted independently. Here we simulate two responses (*K* = 2), *n* = 400, *d* = 5000, the entries of the predictor matrix are i.i.d random signs 1, 1 with probability 0.5 each, and the time-to-event responses are exponential distributed with rate 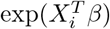, which satisfies proportional hazard. The parameters *β*_1_, *β*_2_ have a common support of size 35. We use a large (5000 samples) and uncensored validation set to select the optimal *λ*. *α* is fixed at 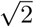. The parameter estimate corresponding to the best *λ* is then evaluated at a large, uncensored test set. We use the C-index as both the validation and test metric. The censoring for both responses are randomly chosen and independent from everything else. The simulations are done for multiple combinations of censoring proportions, each repeated 100 times. Here we report the improvement in C-index of the response with rare events. See Figure 1.

**Figure 1:**
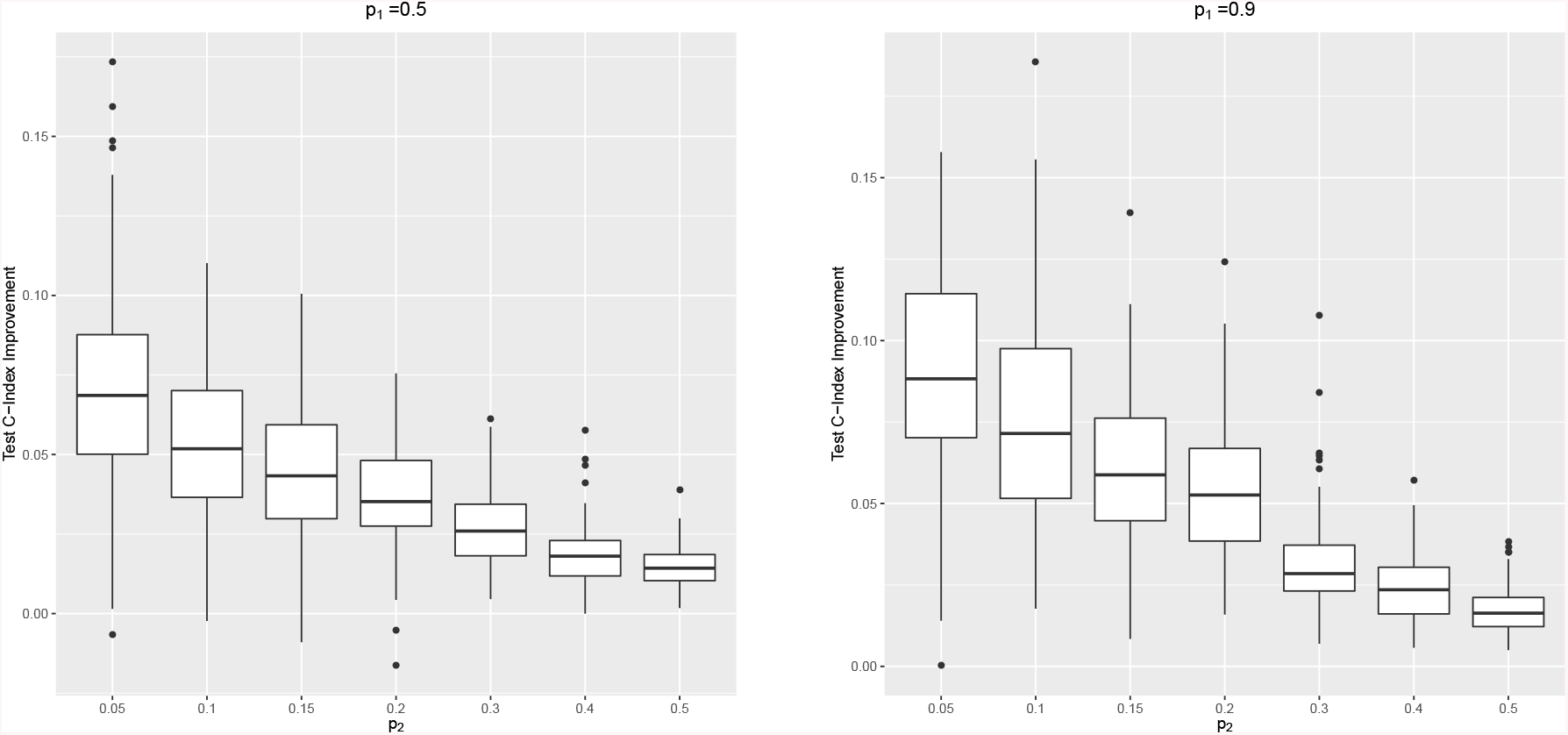
Absolute improvements in test C-index of the rare event response when two responses are trained together. *p*_1_ is the proportion of uncensored events of the first response. *p*_2_ (rare) is the proportion of uncensored events of the second response. The true coefficients have exactly the same support.

The assumption that the coefficients for different responses have the same support can come from domain knowledge (such as biology). In practice this is usually not exactly satisfied. Here we use simulation to examine the robustness of our approach. The setup is the same as the previous simulations, except now the overlap of the support might be smaller than 35 (the number of non-zero entries of *β*_1_, *β*_2_). See the left panel of Figure 2.

**Figure 2:**
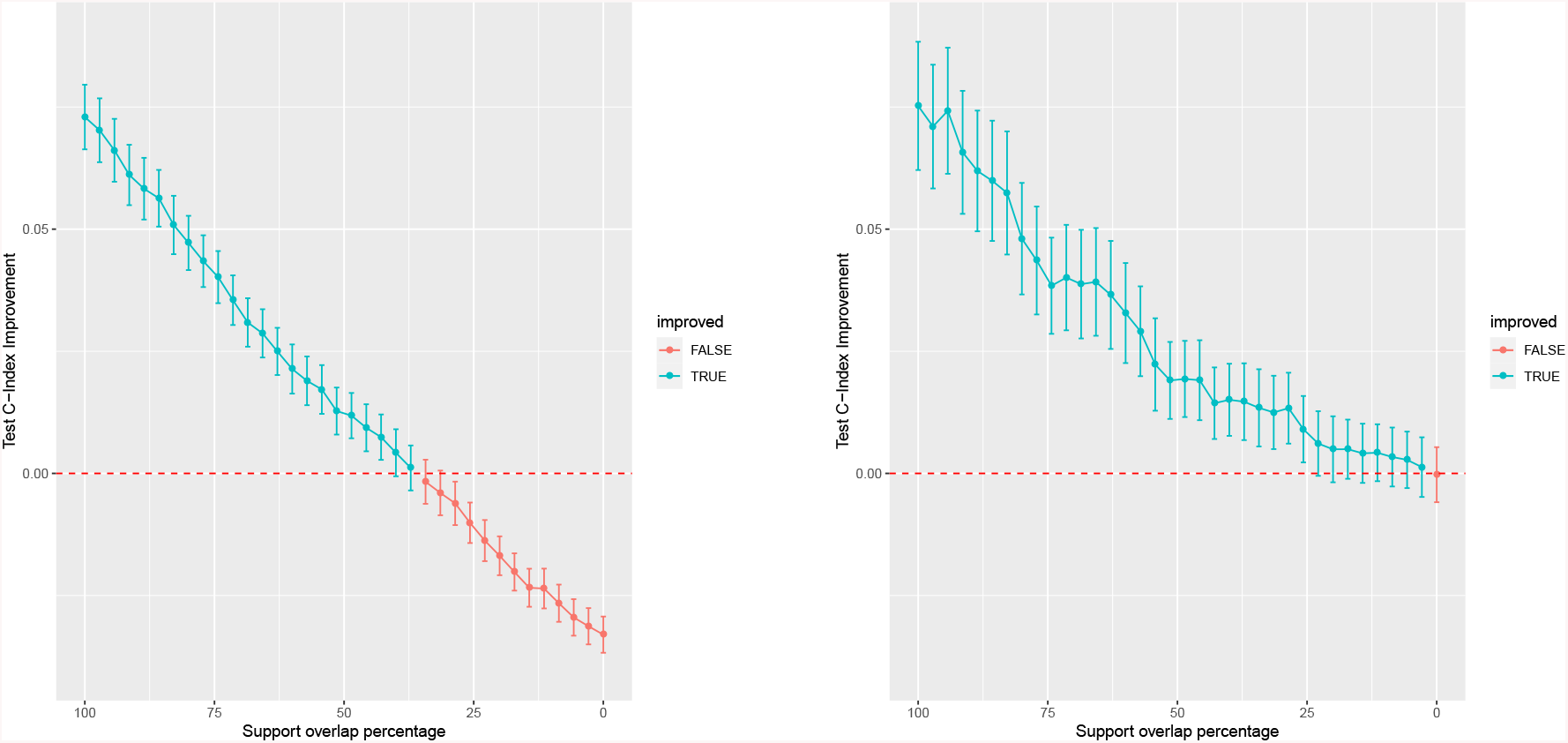
Absolute improvements in Test C-index of the rare event response when two responses are trained together. 60% of the events for the first response are uncensored, and 5% of the events for the second are uncensored. The horizontal axis is the overlap proportion of the support of *β*_1_ and *β*_2_, in other words 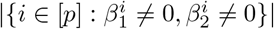. The left panel shows the result when *α* is fixed at 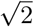, and *λ* is selected from a large uncensored validation set. The right panel shows the result when *α* is selected between 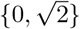 using a small validation set of size 200 with 5% uncensored second event. The whiskers indicates 95% confidence intervals.

We can see that, when the support overlap percentage is less than around 40% the prediction performance actually becomes worse when the two responses are trained together. One solution to this problem is to also treat *α* as a hyperparameter and use the validation set to determine it. For large data set having a two-dimensional hyperparameter could be quite cumbersome. In our simulation and real data application, we only choose *α* from two values 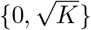, although in principle one can use a large set of *α* candidates at a higher computational cost. The right panel of Figure 2 shows the C-index improvements when we use a small validation set to determine whether *α* should be 0 or 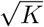. All other settings are the same as above.

## 4 Application to UK Biobank Data

In this section we apply the proposed method to UK Biobank data. We focus on thyroiditis, which has 808 observed events in the study population (337, 129 white British participants). We randomly assign 70% of the samples to the training set, 10% to the validation set, and 20% to the test set. The predictors here are ~ 1 million genetic variants, as well as 11 covariates (sex, and 10 principal components of the genetic variants). We first fit a baseline model using only the 11 covariates, without using any genetic variants. This gives a baseline test C-index at 0.649. We then fit a single-response Lasso Cox regression, where *λ* is selected using based on the validation C-index. At the best *λ* value, the test C-index is 0.679. To apply our approach, we pair thyroiditis with 6 other more common endocrine diseases that we believe share common genetic factors with thyroiditis. These diseases are listed in the table below. Here we report the test C-indices when thyroiditis is paired with one other disease and when all 7 responses are trained together. We also report the test C-indices when we use the validation set to determine both the optimal *λ* and whether to set *α* = 0 or 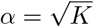.

Table 1 shows clear increase in prediction performance on thyroiditis when all 7 responses are trained together, and when thyroiditis is paired with thyrotoxicosis only. On the other hand, all multi-response solutions have test C-index comparable with the single response solution.

**Table 1:** Test C-index of thyroiditis when this response is paired with other ones. The second column is obtained when *α* is fixed at 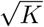 (which is 1 in the first row, 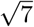 in the last row, and 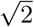 in the rest). The third column is obtained when we use the validation set to choose *α* between 0 (single-response) and 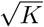. Improved C-indices are given in boldface. Baseline test C-index of thyroiditis, when only the 11 covariates are used for fitting, is 0.649.

| Paired response(s) | Test C-index | Test C-index (validate $\alpha$) |
| --- | --- | --- |
| thyroiditis (single response) | 0.679 | - |
| other hypothyroidism | **0.682** | **0.682** |
| other non-toxic goitre | **0.681** | 0.679 |
| thyrotoxicosis (hyperthyroidism) | **0.686** | **0.686** |
| other disorders of thyroid | 0.679 | 0.679 |
| insulin-dependent diabetes mellitus | **0.683** | **0.683** |
| non-insulin-dependent diabetes mellitus | 0.679 | 0.679 |
| all 7 responses trained together | **0.688** | **0.688** |

## 5 Conclusion and Discussion

We developed a regularized regression method for multiple survival responses. This method is particularly suitable when we can pair a response with few observed events with others having a larger number of events, such that these responses have a same set of useful predictors. We demonstrate the improvements in the prediction accuracy for the rare events responses through simulation and real data applications. We also provide efficient implementation for the proposed method.

Here are two directions for future studies. When there are more than two responses, the relationship between the prediction performance and the degree of overlapping in the coefficients support is yet to be understood. On the practical side, it is reasonable to have different types of models (not just Cox model), or even different response types (such as binary or count) to boost the accuracy of survival analysis on rare events. In principle the same type of regularization could still be applied.

For the GPU implementation, one challenge is the limited amount of memory available on GPUs. In modern computer clusters it is common to have machines with hundreds of Gigabytes of system memory, but most GPUs have less than 32GB of memory. In our implementation we use single precision floating point numbers to store the data matrix, which alleviates memory burden by a factor of two. However, for UK Biobank scale data this is still sometimes insufficient. For example, when the number of participants in the training data is 250, 000, we are only able to fit a model with up to 32, 000 variables on a Tesla V100. One possible solution is to use multiple GPUs, but inter-GPU communication might become the bottleneck. For genetics data, another solution is to utilize their 2-bit representation, which can significantly reduce memory requirement (2 bits per entry vs 32). We leave these ideas for future studies.

## 6 Proof of Propositions

### 6.1 Proof of Proposition 1

We show a slightly more general result from convex analysis. Let *f*: ℝ^*K*^ ⟼ ℝ be continuously differentiable, convex, and bounded from below. Let || · || be a norm on ℝ^*K*^, and || · ||_*_ be its corresponding dual norm. Let *λ* > 0, and set

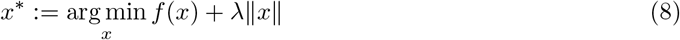

We will show that

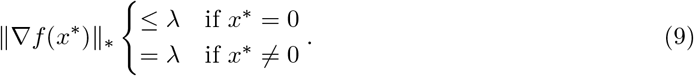

It is clear that (8) is equivalent to the constrained optimization problem

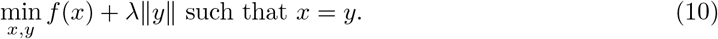

The Lagrangian is

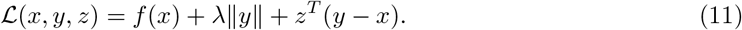

The Lagrangian dual is

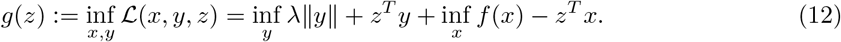

Using the definition of dual norm, when ||*z*||_*\**_ > λ, the infimum of the first term above is −∞, and when ||*z*||_*\**_ ≤ *λ*, the infimum is 0. Therefore

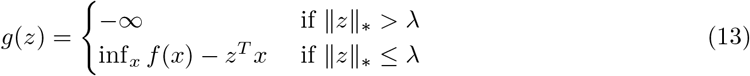

Therefore the dual solution *z*^*^ := arg max_*z*_*g*(*z*) must satisfy ||*z*||_*_ ≦ *λ*. Now we go back to the Lagrangian. Since the primal objective is convex and Slaters condition holds, the solution to the primal problem can be obtained through minimizing

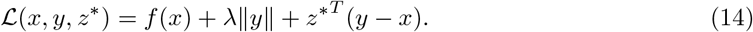

which implies that, at the optimal *x*^*^ we must have ∇*f* (*x*^*\**^) = *z*^*^, so

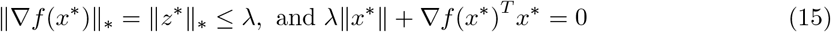

If *x*^*^ = 0, then the second equality is already satisfied, so we only need ||∇*f* (*x*^*^)||_*_ ≤ λ. If *x*^*^ = 0, then by Holder’s inequality

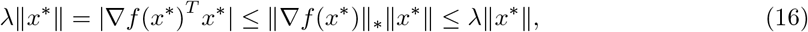

so we must have ||∇*f* (*x*^*^)||_*\**_ = *λ*. Proposition 1 is a direct consequence of this result.

### 6.2 Proof of Proposition 2

We prove the claim for *λ* = 1. The regularization term can be written as

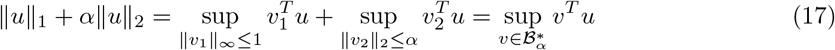

where 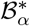 is the unit dual ball

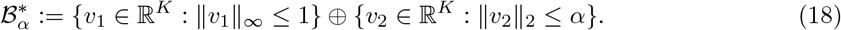

That is ||*v*||_*α*_* ≤ 1 if and only if 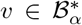, which by definition means that *v* = *v*_1_ + *v*_2_ for some ||*v*_1_||_∞_ ≤ 1, ||*v*_2_||_2_ ≤ *α*. We must have (and it is sufficient to have)

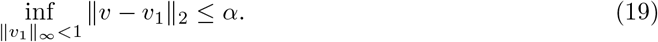

The infimum on the left-hand side is achieved when *v*_1_ = *v* − *S*_1_(*v,* 1), which proves the claim.

## Acknowledgments

Y.T. is supported by Funai Overseas Scholarship from Funai Foundation for Information Technology and the Stanford University School of Medicine.

M.A.R. is supported by Stanford University and a National Institute of Health center for Multi and Trans-ethnic Mapping of Mendelian and Complex Diseases grant (5U01 HG009080). This work was supported by National Human Genome Research Institute (NHGRI) of the National Institutes of Health (NIH) under awards R01HG010140. The content is solely the responsibility of the authors and does not necessarily represent the official views of the National Institutes of Health.

R.T was partially supported by NIH grant 5R01 EB001988-16 and NSF grant 19 DMS1208164.

T.H. was partially supported by grant DMS-1407548 from the National Science Foundation, and grant 5R01 EB 001988-21 from the National Institutes of Health.

This research has been conducted using the UK Biobank Resource under application number 24983. We thank all the participants in the study. The primary and processed data used to generate the analyses presented here are available in the UK Biobank access management system (https://amsportal.ukbiobank.ac.uk/) for application 24983, “Generating effective therapeutic hypotheses from genomic and hospital linkage data” (http://www.ukbiobank.ac.uk/wp-content/uploads/2017/06/24983-Dr-Manuel-Rivas.pdf), and the results are displayed in the Global Biobank Engine (https://biobankengine.stanford.edu).

Some of the computing for this project was performed on the Sherlock cluster. We would like to thank Stanford University and the Stanford Research Computing Center for providing computational resources and support that contributed to these research results.

